# Comparative Genome-Wide Dna Methylation Analysis In Myocardial Tissue From Donors With And Without Down Syndrome

**DOI:** 10.1101/2020.07.15.203075

**Authors:** Romina B. Cejas, Jie Wang, Rachael Hageman-Blair, Song Liu, Javier G. Blanco

## Abstract

Down syndrome (DS, trisomy 21) is the most common major chromosomal aneuploidy compatible with life. The additional whole or partial copy of chromosome 21 results in genome-wide imbalances that drive the complex pathobiology of DS. Differential DNA methylation in the context of trisomy 21 may contribute to the variable architecture of the DS phenotype. The goal of this study was to examine the genomic DNA methylation landscape in myocardial tissue from non-fetal individuals with DS. More than 480,000 unique CpG sites were interrogated in myocardial DNA samples from individuals with (n = 12) and without DS (n = 12) using DNA methylation arrays. A total of 93 highly differentially methylated CpG sites and 16 differentially methylated regions were identified in myocardial DNA from subjects with DS. There were 18 differentially methylated CpG sites in chromosome 21, including 5 highly differentially methylated sites. A CpG site in the *RUNX1* locus was differentially methylated in DS myocardium, and linear regression suggests that donors’ age, gender, DS status, and *RUNX1* methylation may contribute up to ~51% of the variability in *RUNX1* mRNA expression. In DS myocardium, only 58% of the genes overlapping with differentially methylated regions codify for proteins with known functions and 24% are non-coding RNAs. This study provides an initial snapshot on the extent of genome-wide differential methylation in myocardial tissue from persons with DS.

## 1. Introduction

In humans, Down syndrome (DS, trisomy 21) is the most common major chromosomal abnormality compatible with life, with an approximate incidence of 1 in 700 to 1 in 900 live births (1). DS is caused by the presence of a full or partial extra copy of chromosome 21. Individuals with DS exhibit a range of phenotypic characteristics and distinctive physical features. Persons with DS are at increased risk for the development of various comorbidities through the lifespan. In general, the incidence of congenital heart disease, thyroid gland dysfunctions, eye and hearing disorders, pediatric leukemia, and testicular cancer is higher in persons with DS in comparison to individuals without DS (2). For example, the incidence of congenital heart disease in the DS setting is 50 times higher than in the “general population”, and the prevalence of congenital heart defects in individuals with DS ranges from 43% to 58% (3, 4).

Variable DNA methylation levels in specific regions across the genome influences the control of gene expression in different cell types and tissues. In general, relative increases in the levels of methylation in distinct CpG dinucleotides located in gene promoter regions or other regulatory regions may result in transcriptional repression (5). Differential DNA methylation in the context of trisomy 21 may contribute to the complex architecture of the DS phenotype. In this regard, the presence of differentially methylated loci has been examined in some tissues from subjects with DS. For example, Serra-Juhé et al. interrogated approximately 28,000 CpG sites in fetal heart DNA samples from donors with DS and congenital heart defects (CHD) and in samples from donors with DS and without CHD. The authors found that regions in the *GATA4* gene were hypermethylated in cardiac DNA from fetuses with DS with or without CHD, as well as in fetuses with isolated heart malformations (6).

There is a paucity of reports describing the genomic DNA methylation landscape in myocardial tissue from non-fetal individuals with DS. The goal of this pilot study was to examine genome-wide DNA methylation profiles in myocardial DNA samples from donors with and without DS. Myocardial DNA samples were examined to interrogate > 480,000 individual CpG sites using the Illumina HumanMethylation450 BeadChip platform. Next, bioinformatic analysis was performed in order to identify differentially methylated loci in myocardial methylomes. This study contributes new data to continue to define the extent of epigenetic variability in the context of DS.

## 2. Materials and Methods

### 2.1. Human Tissue Specimens

The Institutional Review Board of the State University of New York at Buffalo (UB-IRB) approved this research. UB-IRB determined that this research is not research with human subjects. This research meets exempt criteria, 45 CRF 46,101(b)(4). Human myocardial tissue samples were provided by the National Disease Research Interchange (NDRI), the Cooperative Human Tissue Network (CHTN), and the National Institute of Child Health and Human Development Brain and Tissue Bank (NICHD-BTB). As per Federal and State regulations, tissue banks require every procurement site to obtain informed consent in compliance with all regulations governing that process in writing from any donor of human tissue (or the next of kin thereof) for the use of tissue for research. Each signed consent form is kept on file at the tissue acquisition site and information is never released to any third party. Tissue banks assign computer generated codes to each donor and do not maintain any information that could be used to identify the donor. Tissue banks provide anonymous samples coded with unique sample identification numbers. Procurement protocols for this project were reviewed and approved by NDRI, CHTN, and NICHD-BTB. Myocardium samples from left ventricles weighing 0.8 - 100 g were recovered <20 hours postmortem and snap frozen in liquid nitrogen. Upon arrival samples were stored in liquid nitrogen until further processing. Donor demographics are shown in Supplementary Table 1.

### 2.2. Nucleic Acid Extraction & Illumina Methylation Assay

Genomic DNA from myocardium was extracted and purified with AutoGen QuickGene DNA Tissue Kit S and an AutoGen QuickGene-810 Nucleic Acid Isolation System (AutoGen Inc). DNA concentrations were measured using a Quant-iT PicoGreen dsDNA Assay Kit (Life Technologies, Thermo Fisher Scientific) following the manufacturer’s instructions. The quality of DNA samples (260 nm/230 nm absorbance ratios) was also examined using a NanoDrop 1000 Spectrophotometer (Thermo Scientific). DS status (i.e., trisomy 21) had been previously confirmed by array comparative hybridization as described (7). Genome-wide methylation analysis was performed by interrogating 482,421 unique CpG sites using the Illumina HumanMethylation450 BeadChip platform (Illumina) at the Genomics Shared Resource, Roswell Park Comprehensive Cancer Center, Buffalo, NY (8). BeadChips were scanned using an iScan Reader (Illumina), and analysis was conducted using GenomeStudio v2011.1 (Illumina).

### 2.3. Genome-Wide Methylation Analysis

Genome-wide methylation data, in the form of intensity data (IDAT) files, were compiled in GenomeStudio v2011.1 and processed using the R package minfi following the recommended workflow (9, 10). A detection p-value was generated for every CpG site in each sample to compare the total signal for each probe to the background signal. Samples with high mean detection p-values were defined as poor quality samples and were excluded from downstream analysis. None of the 24 DNA samples were identified as poor-quality sample. Data normalization was performed using the stratified quantile normalization method in the minfi package. Then, probes with unreliable signals were filtered out before differential methylation analysis. Probes with high detection p-value (*p* ≥ 0.01) in any sample were excluded. Probes located on sex chromosomes or those known to have common single nucleotide polymorphisms (SNPs) at the CpG site were filtered out and were not included in subsequent analyses. Probes showing to be cross-reactive due to multiple mapping in the genome were also filtered out (11). All remaining probes were used in the downstream analysis.

Beta values (β value) were used to define methylation, ranging from 0 - 1, for individual probes or genomic regions. Differential methylation analysis was conducted on individual CpG sites, as well as in defined genomic regions (i.e., loci). Individual CpG sites and genomic regions were identified as statistically significantly differentially methylated between DS and non-DS samples following correction for multiple testing using the Benjamini-Hochberg false discovery rate (FDR), with a 5% FDR cut off. Highly differentially methylated CpG sites and regions were identified as those with significant p-values and an absolute methylation difference greater than 0.100 (i.e., β difference > 10.0%). Analysis of differentially methylated regions (DMR) was performed with the DMRcate package, which identifies and ranks the DMR based on a kernel smoothing of the differentially methylated CpG sites (12). Genomic location of individual CpG sites, genic regions, and corresponding gene names were obtained from the UCSC Genome Browser (hg19). Functional classification, and GO enrichment and pathway analyses were performed with the web tool PANTHER (v15.0) (13).

### 2.4. Quantitative Real-time Polymerase Chain Reaction

Total RNA was isolated from myocardium with Trizol reagent according to the manufacturer’s instructions (Thermo Fisher). *RUNX1* mRNA expression was analyzed with specific primers: 5’-CGGTCGAAGTGGAAGAGGGAA-3’ (forward), 5’-ATGGCTCTGTGGTAGGTGGC-3’ (reverse). Total RNA (12.5 ng) was reverse transcribed and amplified with the iTaq Universal SYBR Green One-Step Kit (Bio-Rad). *RUNX1* and *ACTB* were amplified in parallel in a CFX96 Touch Real-Time PCR Detection System (Bio-Rad) with the following cycling parameters: 50°C for 10 min (reverse transcription), 95°C for 1 min, followed by 44 cycles of 95°C for 10 s, 60.5°C for 20 s. Calibration curves were prepared to analyze linearity and PCR efficiency. qRT-PCR data were analyzed using the ΔΔCt method with CFX manager Software (Bio-Rad). The ΔCt method was utilized to determine the relative expression of *RUNX1* mRNA.

### 2.5. Statistical Analyses

For genome-wide methylation analysis, statistical testing was performed with *R* using Fisher tests. Statistical significance was set at FDR of 0.05. β-values were used to represent the extent of methylation at distinct CpG sites. For *RUNX1* mRNA expression analysis, the D’Agostino & Pearson omnibus normality test was used to determine the normality of data sets. Comparisons between means were performed with the Mann-Whitney’s U test.

## 3. Results

### 3.1. DNA methylation profiles in myocardial samples

Data were first preprocessed to remove unreliable probes (1,458 sites), CpG sites containing SNPs (16,928 sites), cross-reactive probes (26,519 sites), and mapping to sex chromosomes (11,295 sites). Following data preprocessing, a total of 429,312 CpG sites were retained for analysis. In general, global methylation profiles of myocardial DNA samples from donors with (n = 12) or without DS (n = 12) were similar; each myocardial DNA methylation profile showed a bimodal distribution of β values, and an average β value of 0.45 (Figure 1A). Donor’s age and gender had no apparent covariate effects on global myocardial DNA methylation profiles.

**Figure 1.**
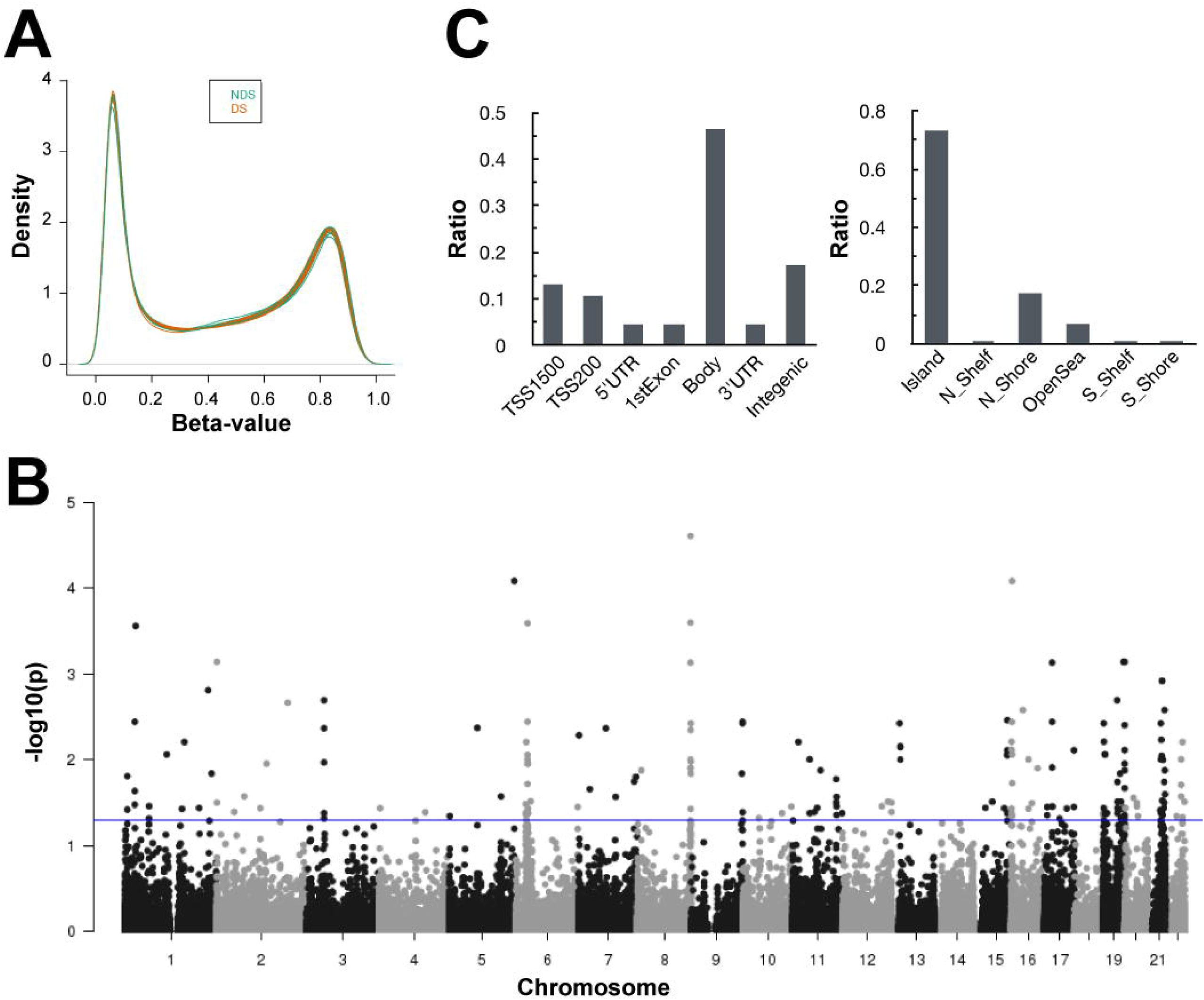
Differential methylation in myocardial DNA from donors with DS. **A)** Density plots of DNA methylation levels (β values) in myocardial DNA from donors with and without DS (DS, n = 12, and non-DS, n = 12). **B)** P values (−log10) of differentially methylated CpG sites stratified by chromosomal location. The blue line depicts the threshold for statistical significance (FDR = 0.05). **C)** Location of differentially methylated sites in relation to functional subregions (left panel) and CpG islands (right panel).

### 3.2. Differentially methylated CpG sites in samples from donors with and without Down syndrome

A total of 192 individual CpG sites were differentially methylated between myocardial DNA samples from subjects with and without DS (FDR < 0.05) (Figure 1B). Of these sites, there were 143 CpG sites with increased methylation in samples with DS relative to samples without DS, whereas 49 CpG sites showed decreased methylation in samples with DS. Analysis of chromosomal locations showed that differentially methylated CpG sites on chromosome 19 were overrepresented and accounted for 15% of the total (29 sites) (Figure 1B). Of all differentially methylated CpG sites, 93 (48%) were observed to be highly differentially methylated (absolute β value difference in DS versus non-DS groups > 0.100) (Table 1). A total of 15 highly differentially methylated CpG sites were found to be less methylated in myocardial DNA samples with DS relative to samples without DS (average β difference = 0.131), whereas 78 CpG sites were more methylated in samples with DS relative to samples without DS (average β difference = 0.145). Of the 93 highly differentially methylated CpG sites, 77 (83%) were distributed in annotated genic regions (i.e., TSS200, TSS1500, 5’ UTR, 1st exon, gene body, 3’ UTR), and 68 (73%) were located within CpG islands (Figure 1C). Genes annotated to differentially methylated CpG sites were functionally classified with the PANTHER Classification System (Table S2). For genes with differentially methylated CpG sites and matching protein products, the majority of the hits (25%) did not match to a specific protein class, and 22% of the listed proteins did not match to a specific molecular function (Figure 2A).

**Table 1.**
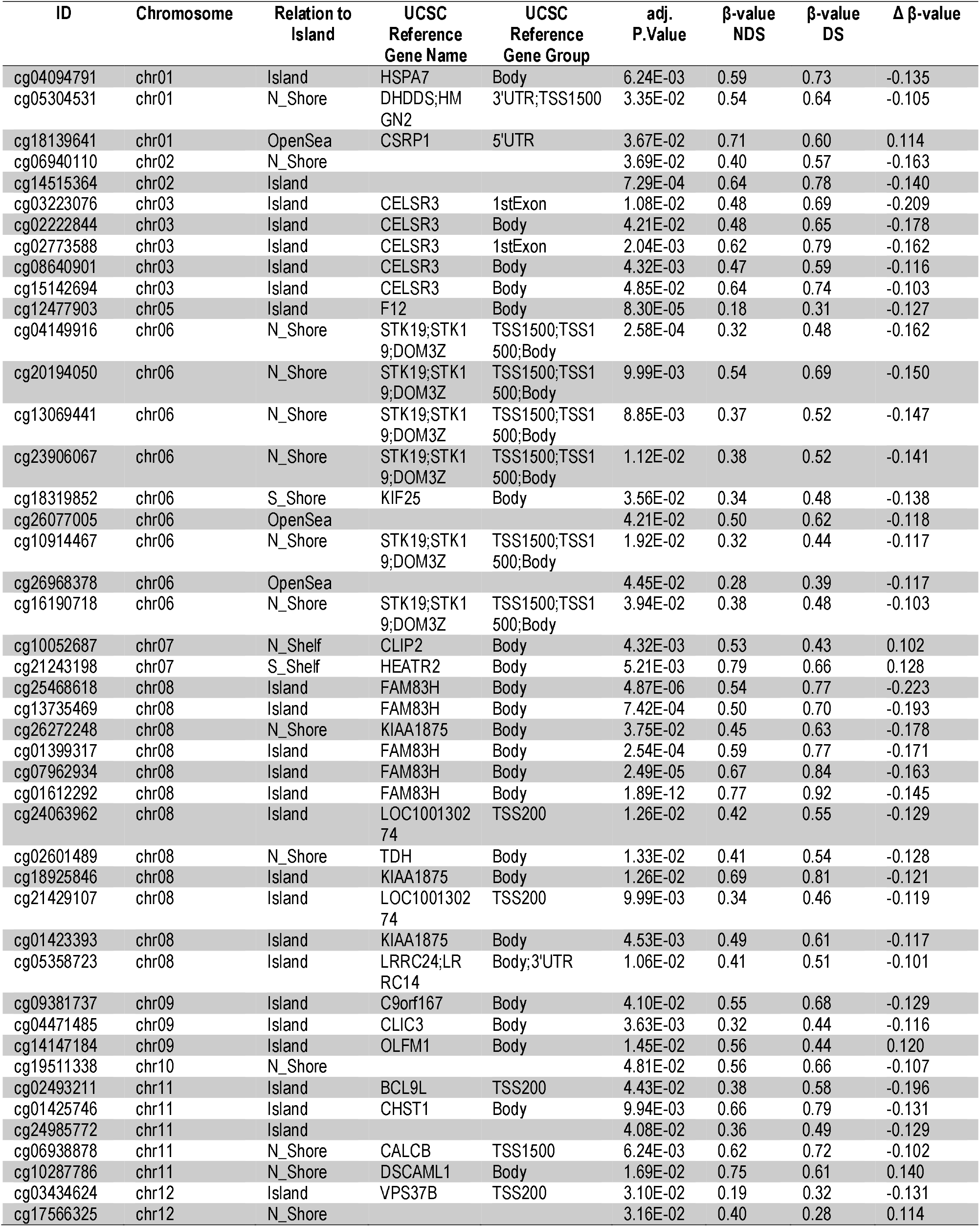
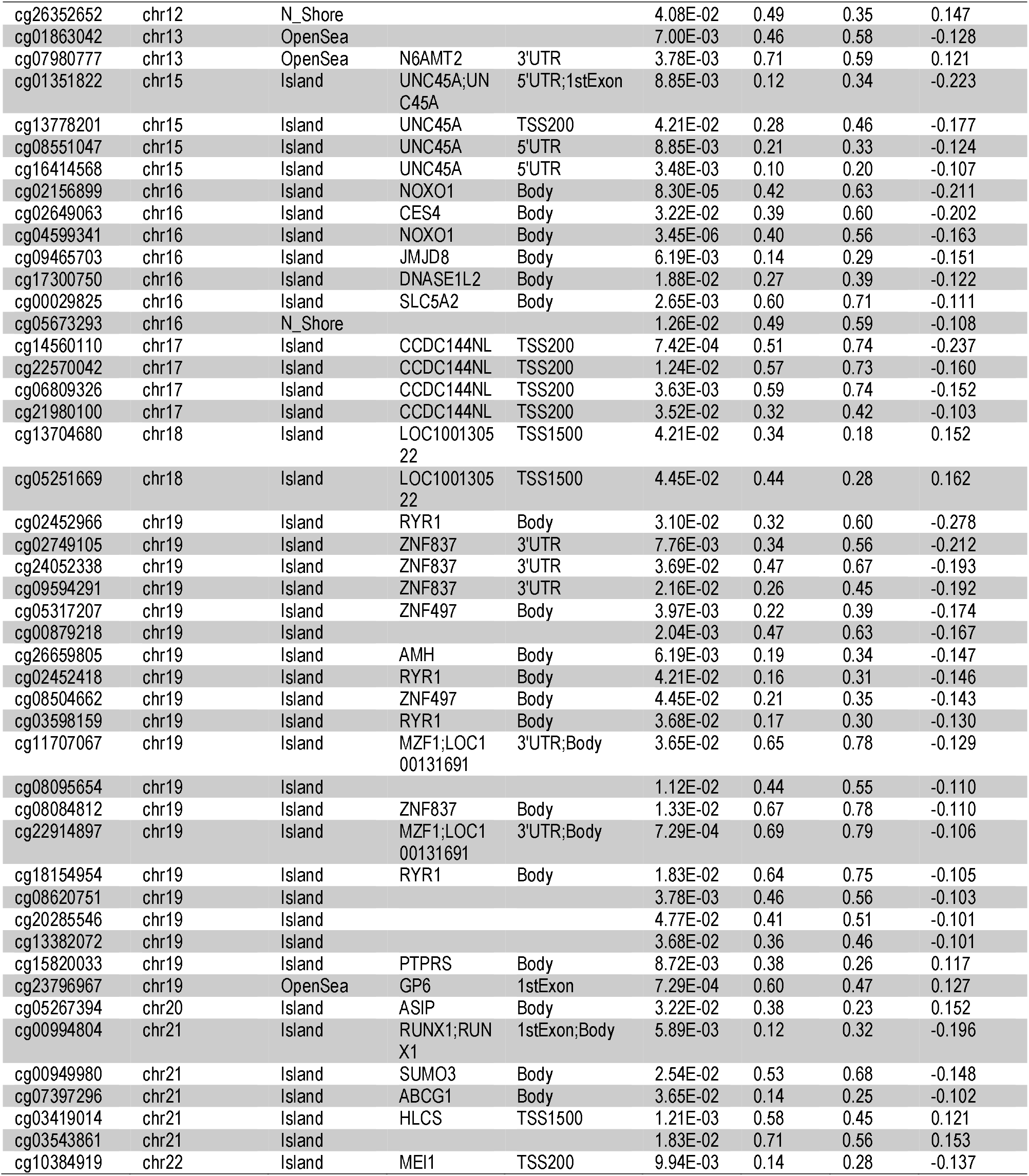
Highly differentially methylated CpG sites in myocardial DNA from subjects with DS.

**Figure 2.**
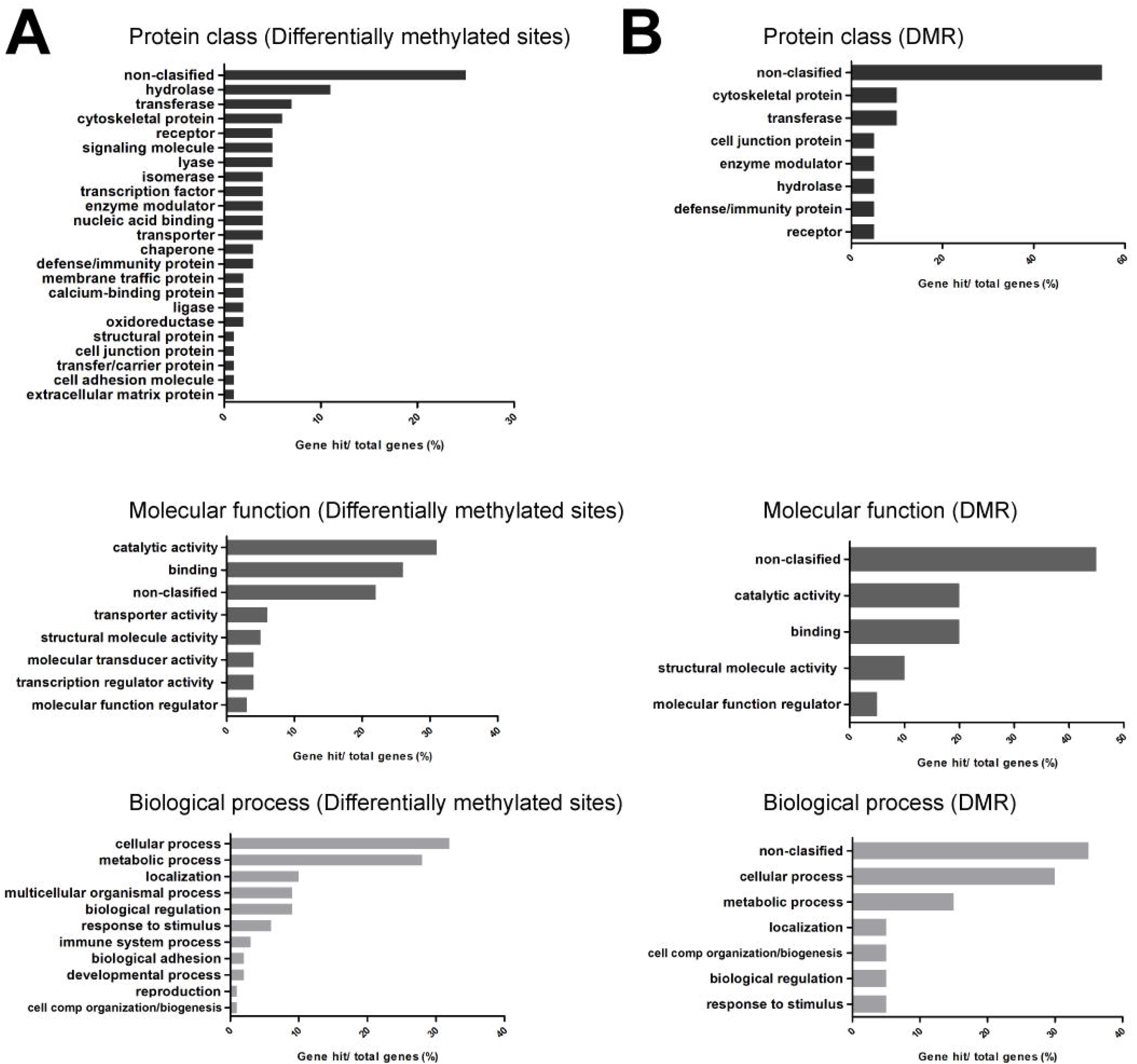
PANTHER functional classification. **A)** Classification of genes mapped to differentially methylated CpG sites. **B)** Classification of genes mapped to differentially methylated regions (DMR). Top panels: protein class, middle panels: molecular function, and bottom panels: biological processes. Genes without hits are listed as non-classified.

### 3.3. Differentially methylated CpG sites on chromosome 21

Special attention was given to differentially methylated CpG sites in chromosome 21 and to sites located in the so-called Down syndrome critical region (DSCR, 21q21.1 – q22.2) (14). Eighteen differentially methylated sites were located in chromosome 21 with a total of 6 highly differentially methylated CpG sites, and 8 differentially methylated CpG sites within the DSCR (Figure 1, and Tables 1 and 2). The mRNA expression of the highly differentially methylated gene *RUNX1* (absolute β value difference = 0.196) was assessed in samples of total myocardial RNA (Table S3). Samples with matching DNA methylation data from donors with DS (n = 8) exhibited higher *RUNX1* mRNA expression (4.96 ± 4.67 relative fold) compared to samples from donors without DS (n = 12, 1.00 ± 0.67 relative fold). The inclusion of additional myocardial mRNA samples with no matching DNA methylation data (non-DS, n = 8, and DS, n = 4) further confirmed the trend towards increased *RUNX1* mRNA expression in DS myocardium (DS: 6.73 ± 6.60 relative fold, and non-DS: 1.00 ± 0.89 relative fold) (Figure 3). Linear regression analysis was performed for the group of myocardial samples with matching *RUNX1* DNA methylation and mRNA expression data (i.e., DS, n = 8, and non-DS, n = 12). Linear regression analysis using donor’s age and gender as covariates suggest that donor status (i.e., DS and non-DS) and *RUNX1* differential methylation expressed as β value contribute to a fraction of the variability observed for the expression of the *RUNX1* transcript in myocardial tissue (R^2^ = 0.61, Adj R^2^ = 0.51, *p* = 0.005). Hypothesis tests on regression coefficients revealed that donor status (DS and non-DS) is significant (*p* = 0.004), although *RUNX1* differential methylation was not found to be significant (*p* = 0.260), possibly due to the small sample size.

**Table 2.**
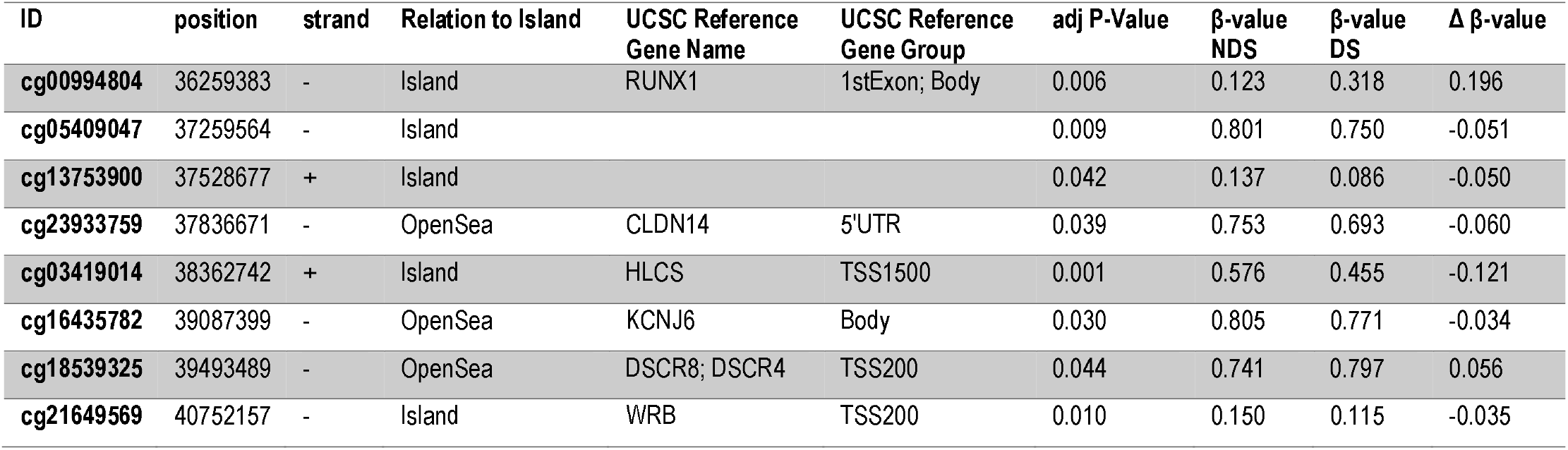
Differentially methylated sites within the DSCR in myocardial DNA from subjects with DS.

**Figure 3.**
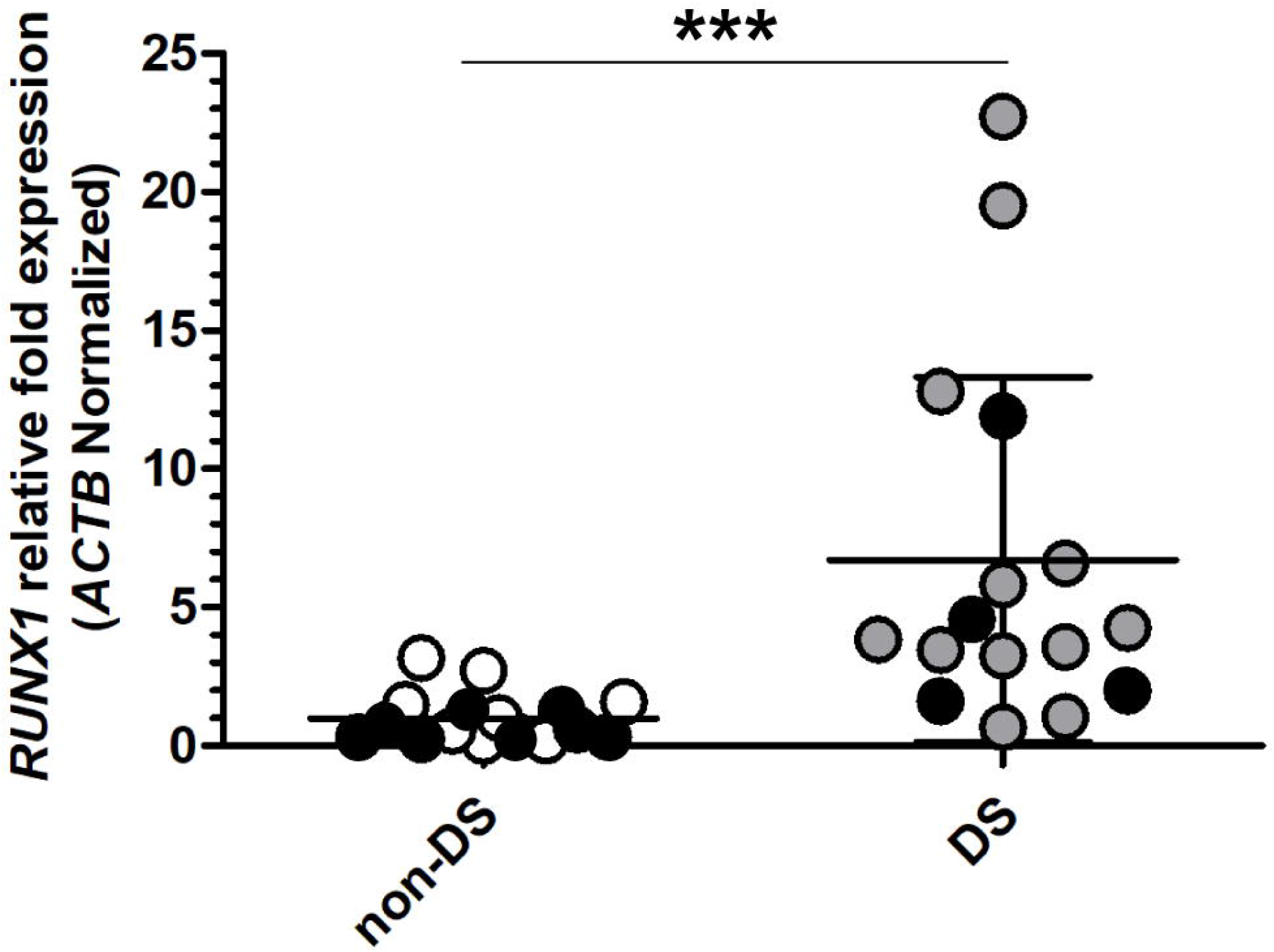
Relative *RUNX1* mRNA fold expression in myocardial samples from donors without and with DS (non-DS, n = 16, and DS, n = 16). Black dots represent additional myocardial mRNA samples from donors without (8 mRNA samples) and with DS (4 mRNA samples) with no matching DNA methylation data. Each point represents the mean from three separate measurements performed in triplicates. Horizontal bars show the mean ± SD. *** P < 0.001, Mann Whitney test.

### 3.4. Differentially methylated regions in DNA samples from donors with and without Down syndrome

A total of 16 differentially methylated regions (DMR) were identified between samples from donors with and without DS (Table 3). There were no DMR in chromosome 21. DMR overlapped with 30 genes, and DMR with the highest number of CpG sites included loci in chromosome 6 (37, 32, 27 and 13 CpG sites, respectively), chromosome 22 (14 CpG sites), and chromosome 18 (10 CpG sites). A total of 20 genes mapped to the PANTHER reference list (Table 4). Genes overlapping with DMR identified by the PANTHER classification system included a relatively high proportion of hits with no specific proteins class (55%), and no assigned molecular function (45%) (Figure 2B). There was no significant enrichment for gene ontology terms (GO) and molecular pathways.

**Table 3.** Differentially methylated regions in myocardial DNA from subjects with DS.

| DMR | Chromosome | Start | End | Width | CpGs # | Min smoothed FDR | Overlapping genes |
| --- | --- | --- | --- | --- | --- | --- | --- |
| 1 | chr06 | 33282313 | 33283849 | 1537 | 37 | 5.24E-87 | SNORA38, ZBTB22, SNORA20 |
| 2 | chr06 | 31938527 | 31939884 | 1358 | 32 | 7.08E-99 | SNORA38, STK19, DXO, SNORA20 |
| 3 | chr06 | 30038754 | 30039380 | 627 | 27 | 1.33E-32 | RNF39, SNORA20 |
| 4 | chr06 | 28829171 | 28829503 | 333 | 13 | 8.22E-33 | RPL13P, XXbac-BPG308K3.6, SNORA20 |
| 5 | chr08 | 144808965 | 144810854 | 1890 | 7 | 5.80E-203 | FAM83H |
| 6 | chr08 | 144790187 | 144790410 | 224 | 5 | 5.70E-32 | ZNF707, CCDC166 |
| 7 | chr13 | 23309774 | 23310464 | 691 | 7 | 1.04E-35 | SNORA16, SNORD37 |
| 8 | chr15 | 91473059 | 91473569 | 511 | 9 | 2.78E-51 | UNC45A |
| 9 | chr16 | 2029256 | 2030009 | 754 | 4 | 4.15E-59 | TBL3, NOXO1 |
| 10 | chr17 | 20799408 | 20799644 | 237 | 6 | 4.83E-43 | SNORA69, RP11-344E13.3, CCDC144NL |
| 11 | chr18 | 77905119 | 77905947 | 829 | 10 | 2.57E-35 | ADNP2, AC139100.2 |
| 12 | chr19 | 55549414 | 55549842 | 429 | 8 | 4.79E-35 | CTC-550B14.7, GP6, CTC-550B14.6 |
| 13 | chr19 | 2251588 | 2252029 | 442 | 4 | 1.53E-35 | AMH |
| 14 | chr22 | 45809244 | 45809793 | 550 | 14 | 4.71E-43 | RIBC2, SMC1B |
| 15 | chr22 | 42095347 | 42095536 | 190 | 6 | 5.52E-32 | MEI1 |
| 16 | chr22 | 51016703 | 51016950 | 248 | 3 | 1.64E-30 | CPT1B, CHKB-CPT1B |

**Table 4.** Genes showing overlap with DMR in myocardial DNA from subjects with DS. Functional classification according to the PANTHER classification system.

| Gene ID | Gene name | GO database Biological Process Complete |
| --- | --- | --- |
| AC139100 | Partitioning defective 6 homolog gamma | centrosome cycle; establishment or maintenance of cell polarity; cell division; regulation of cellular localization; bicellular tight junction assembly |
| ADNP2 | Activity-dependent neuroprotector homeobox protein 2 | regulation of transcription by RNA polymerase II; neuron differentiation; positive regulation of cell growth; cellular response to oxidative stress; negative regulation of cell death; cellular response to retinoic acid |
| AMH | Muellerian-inhibiting factor | preantral ovarian follicle growth; urogenital system development; Mullerian duct regression; cell-cell signaling; gonadal mesoderm development; sex determination; sex differentiation; aging; positive regulation of gene expression; response to organic cyclic compound; BMP signaling pathway; response to drug; positive regulation of NF-kappaB transcription factor activity; negative regulation of ovarian follicle development |
| CCDC144NL | Putative coiled-coil domain-containing protein 144 N-terminal-like |  |
| CCDC166 | Coiled-coil domain-containing protein 166 |  |
| CHKB-CPT1B | CHKB-CPT1B readthrough (NMD candidate) (Fragment) | phosphorylation; glycerophospholipid biosynthetic process |
| CPT1B | Carnitine O-palmitoyltransferase 1, muscle isoform | fatty acid metabolic process; fatty acid beta-oxidation; carnitine shuttle; carnitine metabolic process; long-chain fatty acid transport |
| DXO | Decapping and exoribonuclease protein | mRNA catabolic process; RNA destabilization; nuclear mRNA surveillance; nucleic acid phosphodiester bond hydrolysis; NAD-cap decapping |
| FAM83H | Protein FAM83H | positive regulation of cell migration; biomineral tissue development; protein localization to cytoskeleton; intermediate filament cytoskeleton organization |
| GP6 | Platelet glycoprotein VI | enzyme linked receptor protein signaling pathway; blood coagulation; platelet activation; leukocyte migration |
| MEI1 | Meiosis inhibitor protein 1 | meiosis I; male meiosis I; spermatid development; meiotic telomere clustering |
| NOXO1 | NADPH oxidase organizer 1 | superoxide metabolic process; regulation of hydrogen peroxide metabolic process; extracellular matrix disassembly; positive regulation of catalytic activity; regulation of respiratory burst |
| RIBC2 | RIB43A-like with coiled-coils protein 2 |  |
| RNF39 | RING finger protein 39 | biological_process |
| SMC1B | Structural maintenance of chromosomes protein 1B | sister chromatid cohesion; meiotic cell cycle |
| STK19 | Serine/threonine-protein kinase 19 | protein phosphorylation; phosphorylation; positive regulation of Ras protein signal transduction |
| TBL3 | Transducin beta-like protein 3 | maturation of SSU-rRNA from tricistronic rRNA transcript (SSU-rRNA, 5.8S rRNA, LSU-rRNA; rRNA processing |
| UNC45A | Protein unc-45 homolog A | muscle organ development; cell differentiation; chaperone-mediated protein folding |
| ZBTB22 | Zinc finger and BTB domain-containing protein 22 | regulation of transcription by RNA polymerase II |
| ZNF707 | Zinc finger protein 707 | regulation of transcription, DNA-templated |
| CTC-550B14.6 | Unable to map |  |
| CTC-550B14.7 | Unable to map |  |
| RP11-344E13.3 | Unable to map |  |
| RPL13P | Unable to map |  |
| SNORA16 | Unable to map |  |
| SNORA20 | Unable to map |  |
| SNORA38 | Unable to map |  |
| SNORA69 | Unable to map |  |
| SNORD37 | Unable to map |  |

## 4. Discussion and Conclusions

This pilot study documents the extent of genome-wide DNA methylation in a collection of samples of myocardial tissue from individuals with and without DS. Persons with DS are at increased risk for the development of various comorbidities which may impact quality of life and mortality rate. The estimated number of people with DS living in the United States was 1 per 1,499 inhabitants in the year 2010 (15). The life expectancy for persons with DS has increased from 9 - 11 years of age in the 1900s to over 55 - 60 years of age in the 2000s, with around 10% of individuals living now up to 70 years of age (16, 17). The myocardial samples analyzed in this study have been obtained from donors representing almost the entire life span of individuals with DS (Table S1). Although donor’s age and gender may potentially impact tissular DNA methylation patterns, it was not possible to fully match both sets of samples by age because the DS group included three samples from young donors (ages: 1, 1, and 10 years old) for which there were not age-matched samples from donors without DS (18–20). Nevertheless, the myocardial samples from young donors with DS were maintained for analysis due to the general scarcity of epigenetic data derived from individuals with DS.

Sailani et al. applied reduced representation bisulfite sequencing (RRBS) for the analysis of methylation patterns in DNA from skin fibroblasts from monozygotic twins with and without trisomy 21. The authors found 35 differentially methylated regions (absolute methylation differences > 25%) with enrichment in the promoter regions of genes involved in embryonic organ morphogenesis (e.g., *HOXB5*, *HOXB6*, *HOXD3*, and *HOXD10)*. The authors also reported an increase in global DNA methylation in trisomic methylomes in comparison with euploid methylomes (21). RRBS of DNA from placental villi from euploid and trisomic pregnancies showed global hypermethylation in samples with trisomy 21 (22). Our comparisons revealed that global DNA methylation levels are similar in myocardial samples from subjects with and without DS (Figure 1). We identified a relatively small number of differentially methylated CpG sites and regions in the myocardial methylomes from donors with DS (Figure 1, and Tables 1 and 3). Thus, it appears that myocardial DNA methylation profiles in the DS context exhibit subtle changes, and differences in DNA methylation levels relative to the non-DS counterpart tend to be localized in a relatively small number of loci. Differentially methylated loci showed no enrichment for GO terms or distinct pathways. In terms of relative methylation at individual CpG sites, there was a trend towards increased methylation in the DS context with 65.7% of total differentially methylated CpG sites and 83.9% of highly differentially methylated CpG sites showing increased methylation in DS samples in comparison to non-DS samples.

Our analysis identified differential methylation in CpG sites and regions overlapping genes with demonstrated roles in cardiac structure and function (i.e., *UNC45A*, *DNM2*), atrioventricular valve formation (i.e., *OLFM1*), and other cardiac processes including calcium homeostasis, muscle contraction, and cardiac conduction (i.e., *DMPK, RYR1*) (Table 4 and Table S2) (23–27). Previous studies have noted differential methylation in CpG sites within some of the loci identified in our analysis. For example, differential methylation in *KIAA1875, CELSR3*, *STK19* and *UNC45A* has been detected in DNA from epithelial cells, placenta, and fetal cortex with trisomy 21 (22, 28, 29). Likewise, we identified a number of highly differentially methylated sites located within the *KIAA1875*, *CELSR3*, *STK19*, and *UNC45A* loci (3, 5, 6, and 4 CpG sites, respectively), all of them showing increased methylation in myocardial DNA from donors with DS (Table 1).

Our study identified 8 CpG sites within the so called DSCR which were differentially methylated in DS myocardium. The DSCR included a highly differentially methylated CpG site within the *RUNX1* locus (Table 2). On average, *RUNX1* mRNA expression in trisomic myocardium was ~6-fold higher than in diploid tissue, although there was considerable interindividual variability (SD: ~6-fold) (Figure 3). RUNX family members form heterodimeric transcription factors that activate or repress gene transcription in healthy and disease states. Little is known regarding the roles of RUNX factors during cardiac physiology. A recent study in a murine model showed that Runx1 modulates calcium ion uptake in sarcoplasmic reticulum and cardiac contractility. Of note, reduction of Runx1 function prevented adverse cardiac remodeling after myocardial infarction (30). Our simple linear regression model based on donors’ age, gender, DS status, and *RUNX1* methylation suggests that these variables may contribute up to ~51% of the variability in myocardial *RUNX1* mRNA expression. Although DS status was found to be a strong predictor of *RUNX1* mRNA methylation, *RUNX1* interindividual methylation did not reach the level of significance *per se*. We hypothesize that this is due in part to the small sample size, which is a limitation for the linear regression. Elucidation of the role of differential *RUNX1* methylation during myocardial function in the context of DS deserves further investigation.

Many of the “genes” overlapping with differentially methylated CpG sites and DMR had unknown function and/or protein-coding potential, or codify non-protein products (Figure 2, Table 4). Only 58% of the genes overlapping with DMR codify for proteins with known functions, and around 24% codify non-coding RNAs. Differential expression of non-coding RNA species has been detected in human trisomic endothelial progenitor cells and induced pluripotent stem cells (31, 32). Many non-coding RNAs regulate complex biological functions and are involved in disease processes such as cardiovascular disease (33, 34). For example, our data suggest that *RNARP11-344E13.3* is differentially methylated in DS myocardium. *RNARP11-344E13.3* is a long non-coding RNA that interacts with microRNA targets linked to the development of cardiac hypertrophy (35). It would be of interest to further investigate the role of differential DNA methylation during the expression of non-coding RNAs in myocardium with emphasis on the DS setting.

This pilot study is limited by the small number of myocardial samples from donors with and without DS. Although our analysis identified a number of differentially methylated loci in DS myocardium, these observations need to be confirmed in larger sample sizes. Research on fundamental issues concerning the pathobiology of DS continues to be hampered by the scarcity of good quality tissue samples from donors with DS. We believe that this study provides an initial snapshot on the extent of genome-wide differential methylation in myocardial tissue from persons with DS. This study contributes new data to further examine the contribution of differential DNA methylation to the complex pathobiology of DS.

## Abbreviations

DS: Down syndrome
non-DS: non-Down syndrome
DNA: deoxyribonucleic acid
*RUNX1*: runt-related transcription factor 1
RNA: ribonucleic acid
mRNA: messenger RNA
DMR: differentially methylated region
SNP: single nucleotide polymorphism
FDR: false discovery rate
*ACTB*: actin beta
PCR: polymerase chain reaction
RT-qPCR: quantitative reverse transcription polymerase chain reaction
DSCR: Down syndrome critical region
GO: gene ontology
RRBS: reduced representation bisulfite sequencing.

