## Supplementary material for "Comparative Genome-Wide Dna Methylation Analysis In Myocardial Tissue From Donors With And Without Down Syndrome"

**Supplementary Table 1.** Donor demographics

| Sample ID | Sample Group | Gender | Age (years) | Relevant diagnoses and pathophysiology |
|---|---|---|---|---|
| **Heart_1** | NDS | Male | 57 | expired d/t intracranial hemorrhage, obese |
| **Heart_5** | NDS | Male | 61 | expired d/t cardiopulmonary arrest |
| **Heart_9** | NDS | Male | 70 | expired d/t cardiac arrest, history of mononeuritis/COPD |
| **Heart_14** | NDS | Male | 80 | expired d/t respiratory failure, history of distant colon cancer, COPD, pulmonary fibrosis |
| **Heart_21** | NDS | Female | 47 | Expired d/t complications with liver, liver transplant secondary to chirrosis. On sirolimus and steroids |
| **Heart_24** | NDS | Male | 69 | Expired d/t complications associated with abdominal aortic aneurysm surgery, probable sepsis secondary to catheter. |
| **Heart_35** | NDS | Male | 42 | expired d/t complications of kidney and pancreas transplant. |
| **Heart_36** | NDS | Male | 68 | expired d/t renal failure. had CHF/CAD |
| **Heart_43** | NDS | Male | 42 | expired d/t ventricular fibrilation, CAD, myocardia ishemia, myocardial infarction. |
| **Heart_53** | NDS | Female | 38 | Obese, hypothiroidism |
| **Heart_54** | NDS | Male | 47 | Hypertension, hepatitis, |
| **Heart_46** | NDS | Female | 71 | expired d/t multiorgan failure and CHF. Mild atherosclerosis, multiple myleoma, on thalidomide and cyclophosphamide. |
| **Heart_18** | DS | Male | 58 | expired d/t respiratory arrest, had pneumonia |
| **Heart_17** | DS | Male | 63 | expired d/t respiratory arrest secondary to subdural hematoma, history of Alzheimer's with dimentia |
| **Heart _15** | DS | Male | 72 | expired d/t aspiration pneumonia. |
| **Heart_37** | DS | Male | 22 | expired spontaneously, hx of heart murmor |
| **Heart_48** | DS | Female | 48 | expired d/t trauma |
| **Heart_23** | DS | Male | 59 | expired d/t coronary arrest, history of CKD/wound infections |
| **Heart_39** | DS | Female | 1 | expired d/t unknown reasons spontaneously. Had upper respiratory infection |
| **Heart_19** | DS | Male | 39 | expired d/t pneumonia, no positive blood cultures, presumed negative for sepsis |
| **Heart_41** | DS | Male | 64 | expired d/t respiratory failure and CVA. |
| **Heart_33** | DS | Male | 40 | expired d/t stage IV adenocarcinoma (no chemo given) and MRSA pneumonia. |
| **Heart_38** | DS | Male | 1 | expired d/t unknown reasons spontaneously. |
| **Heart_40** | DS | Male | 10 | expired d/t cardiac arrest, unremarkable history |

**Supplementary Table 2.** Genes showing overlap with differentially methylated sites in myocardial DNA from subjects with DS. Functional classification according to the PANTHER classification system.

| Gene ID | Gene name | GO database Biological Process Complete |
|---|---|---|
| **ABCG1** | ATP-binding cassette sub-family G member 1 | cholesterol metabolic process(GO:0008203);response to organic substance(GO:0010033);negative regulation of macrophage derived foam cell differentiation(GO:0010745);regulation of cholesterol esterification(GO:0010872);positive regulation of cholesterol efflux(GO:0010875);negative regulation of cholesterol storage(GO:0010887);intracellular receptor signaling pathway(GO:0030522);intracellular cholesterol transport(GO:0032367);cholesterol efflux(GO:0033344);phospholipid efflux(GO:0033700);response to lipid(GO:0033993);low-density lipoprotein particle remodeling(GO:0034374);high-density lipoprotein particle remodeling(GO:0034375);glycoprotein transport(GO:0034436);cholesterol homeostasis(GO:0042632);amyloid precursor protein catabolic process(GO:0042987);reverse cholesterol transport(GO:0043691);positive regulation of cholesterol biosynthetic process(GO:0045542);positive regulation of protein secretion(GO:0050714);transmembrane transport(GO:0055085);phospholipid homeostasis(GO:0055091);cellular response to high density lipoprotein particle stimulus(GO:0071403);toxin transport(GO:1901998);positive regulation of amyloid-beta formation(GO:1902004) |
| **ACAP1** | Arf-GAP with coiled-coil, ANK repeat and PH domain-containing protein 1 | protein transport(GO:0015031);positive regulation of GTPase activity(GO:0043547) |
| **ACMSD** | 2-amino-3-carboxymuconate-6-semialdehyde decarboxylase | tryptophan catabolic process(GO:0006569);secondary metabolic process(GO:0019748);negative regulation of quinolinate biosynthetic process(GO:1904985);picolinic acid biosynthetic process(GO:1905004);regulation of 'de novo' NAD biosynthetic process from tryptophan(GO:1905012) |
| **AMH** | Muellerian-inhibiting factor | preantral ovarian follicle growth(GO:0001546);urogenital system development(GO:0001655);Mullerian duct regression(GO:0001880);cell-cell signaling(GO:0007267);gonadal mesoderm development(GO:0007506);sex determination(GO:0007530);sex differentiation(GO:0007548);aging(GO:0007568);positive regulation of gene expression(GO:0010628);response to organic cyclic compound(GO:0014070);BMP signaling pathway(GO:0030509);response to drug(GO:0042493);positive regulation of NF-kappaB transcription factor activity(GO:0051092);negative regulation of ovarian follicle development(GO:2000355) |
| **ANKRD26P1** | Putative ankyrin repeat domain-containing protein 26-like protein | |
| **AP3B2** | AP-3 complex subunit beta-2 | intracellular protein transport(GO:0006886);anterograde axonal transport(GO:0008089);protein transport(GO:0015031);vesicle-mediated transport(GO:0016192);anterograde synaptic vesicle transport(GO:0048490) |
| **ASIP** | Agouti-signaling protein | generation of precursor metabolites and energy(GO:0006091);signal transduction(GO:0007165);cell-cell signaling(GO:0007267);adult feeding behavior(GO:0008343);hormone-mediated signaling pathway(GO:0009755);melanosome transport(GO:0032402);melanosome organization(GO:0032438);regulation of molecular function, epigenetic(GO:0040030);melanin biosynthetic process(GO:0042438);positive regulation of melanin biosynthetic process(GO:0048023);genetic imprinting(GO:0071514) |
| **BCL9L** | B-cell CLL/lymphoma 9-like protein | positive regulation of epithelial to mesenchymal transition(GO:0010718);regulation of cell morphogenesis(GO:0022604);negative regulation of transforming growth factor beta receptor signaling pathway(GO:0030512);somatic stem cell population maintenance(GO:0035019);skeletal muscle cell differentiation(GO:0035914);positive regulation of transcription by RNA polymerase II(GO:0045944);canonical Wnt signaling pathway(GO:0060070);positive regulation of nucleic acid-templated transcription(GO:1903508);beta-catenin-TCF complex assembly(GO:1904837) |
| **C17orf90** | Oxidoreductase-like domain-containing protein 1 | |
| **C8ORFK29** | Transmembrane protein 249 | |
| **C9orf167** | Torsin-4A | platelet degranulation(GO:0002576) |
| **CALCB** | Calcitonin gene-related peptide 2 | cellular calcium ion homeostasis(GO:0006874);signal transduction(GO:0007165);G protein-coupled receptor signaling pathway(GO:0007186);adenylate cyclase-activating G protein-coupled receptor signaling pathway(GO:0007189);regulation of cytosolic calcium ion concentration(GO:0051480) |
| **CBS** | Cystathionine beta-synthase | cysteine biosynthetic process from serine(GO:0006535);L-serine metabolic process(GO:0006563);L-serine catabolic process(GO:0006565);cysteine biosynthetic process via cystathionine(GO:0019343);cysteine biosynthetic process(GO:0019344);transsulfuration(GO:0019346);L-cysteine catabolic process(GO:0019448);DNA protection(GO:0042262);homocysteine catabolic process(GO:0043418);homocysteine metabolic process(GO:0050667);oxidation-reduction process(GO:0055114);hydrogen sulfide biosynthetic process(GO:0070814) |
| **CCDC144NL** | Putative coiled-coil domain-containing protein 144 N-terminal-like | |
| **CELSR3** | Cadherin EGF LAG seven-pass G-type receptor 3 | neuron migration(GO:0001764);regulation of protein phosphorylation(GO:0001932);homophilic cell adhesion via plasma membrane adhesion molecules(GO:0007156);G protein-coupled receptor signaling pathway(GO:0007186);axonal fasciculation(GO:0007413);regulation of protein localization(GO:0032880);dopaminergic neuron axon guidance(GO:0036514);serotonergic neuron axon guidance(GO:0036515);Wnt signaling pathway, planar cell polarity pathway(GO:0060071);cilium assembly(GO:0060271);cell-cell adhesion(GO:0098609);planar cell polarity pathway involved in axon guidance(GO:1904938) |
| **CES4** | Putative inactive carboxylesterase 4 | anatomical structure morphogenesis(GO:0009653);lipid catabolic process(GO:0016042) |
| **CHST1** | Carbohydrate sulfotransferase 1 | polysaccharide metabolic process(GO:0005976);galactose metabolic process(GO:0006012);N-acetylglucosamine metabolic process(GO:0006044);sulfur compound metabolic process(GO:0006790);inflammatory response(GO:0006954);keratan sulfate biosynthetic process(GO:0018146);keratan sulfate metabolic process(GO:0042339) |
| **CLDN14** | Claudin-14 | calcium-independent cell-cell adhesion via plasma membrane cell-adhesion molecules(GO:0016338);protein-containing complex assembly(GO:0065003) |
| **CLIC3** | Chloride intracellular channel protein 3 | chloride transport(GO:0006821);signal transduction(GO:0007165);regulation of ion transmembrane transport(GO:0034765);chloride transmembrane transport(GO:1902476) |
| **CLIP2** | CAP-Gly domain-containing linker protein 2 | |
| **CLPTM1L** | Cleft lip and palate transmembrane protein 1-like protein | apoptotic process(GO:0006915) |
| **CRHBP** | Corticotropin-releasing factor-binding protein | synaptic transmission, dopaminergic(GO:0001963);maternal aggressive behavior(GO:0002125);inflammatory response(GO:0006954);signal transduction(GO:0007165);female pregnancy(GO:0007565);learning or |

| | | |
|---|---|---|
| | | memory(GO:0007611);hormone-mediated signaling pathway(GO:0009755);cellular response to drug(GO:0035690);cellular response to potassium ion(GO:0035865);hormone metabolic process(GO:0042445);regulated exocytosis(GO:0045055);behavioral response to ethanol(GO:0048149);regulation of corticotropin secretion(GO:0051459);negative regulation of corticotropin secretion(GO:0051460);cellular response to calcium ion(GO:0071277);cellular response to cocaine(GO:0071314);cellular response to cAMP(GO:0071320);cellular response to tumor necrosis factor(GO:0071356);cellular response to estrogen stimulus(GO:0071391);cellular response to estradiol stimulus(GO:0071392);regulation of cellular response to stress(GO:0080135);cellular response to gonadotropin-releasing hormone(GO:0097211);negative regulation of corticotropin-releasing hormone receptor activity(GO:1900011);regulation of NMDA receptor activity(GO:2000310) |
| **CRYZL1** | Quinone oxidoreductase-like protein 1 | oxidation-reduction process(GO:0055114);quinone metabolic process(GO:1901661) |
| **CSRP1** | Cysteine and glycine-rich protein 1 | actin cytoskeleton organization(GO:0030036);sarcomere organization(GO:0045214);muscle tissue development(GO:0060537);platelet aggregation(GO:0070527) |
| **CYTH2** | Cytohesin-2 | endocytosis(GO:0006897);actin cytoskeleton organization(GO:0030036);regulation of ARF protein signal transduction(GO:0032012) |
| **DCXR** | L-xylulose reductase | xylulose metabolic process(GO:0005997);glucose metabolic process(GO:0006006);NADP metabolic process(GO:0006739);glucuronate catabolic process to xylulose 5-phosphate(GO:0019640);D-xylose metabolic process(GO:0042732);oxidation-reduction process(GO:0055114) |
| **DHDDS** | Dehydrodolichyl diphosphate synthase complex subunit DHDDS | dolichyl diphosphate biosynthetic process(GO:0006489);polyprenol biosynthetic process(GO:0016094) |
| **DMPK** | Myotonin-protein kinase | regulation of sodium ion transport(GO:0002028);protein phosphorylation(GO:0006468);cellular calcium ion homeostasis(GO:0006874);nuclear envelope organization(GO:0006998);regulation of heart contraction(GO:0008016);muscle cell apoptotic process(GO:0010657);regulation of myotube differentiation(GO:0010830);regulation of skeletal muscle contraction by calcium ion signaling(GO:0014722);regulation of excitatory postsynaptic membrane potential involved in skeletal muscle contraction(GO:0014853);peptidyl-serine phosphorylation(GO:0018105);intracellular signal transduction(GO:0035556);regulation of phosphoprotein phosphatase activity(GO:0043666);regulation of synapse structural plasticity(GO:0051823);regulation of cardiac conduction(GO:1903779) |
| **DNAAF5** | Dynein assembly factor 5, axonemal | cilium movement(GO:0003341);outer dynein arm assembly(GO:0036158);inner dynein arm assembly(GO:0036159) |
| **DNAJC6** | Putative tyrosine-protein phosphatase auxilin | post-Golgi vesicle-mediated transport(GO:0006892);receptor-mediated endocytosis(GO:0006898);synaptic vesicle uncoating(GO:0016191);peptidyl-tyrosine dephosphorylation(GO:0035335);membrane organization(GO:0061024);clathrin coat disassembly(GO:0072318);clathrin-dependent endocytosis(GO:0072583);regulation of clathrin-dependent endocytosis(GO:2000369) |
| **DNASE1L2** | Deoxyribonuclease-1-like 2 | DNA catabolic process, endonucleolytic(GO:0000737);hair follicle development(GO:0001942);corneocyte development(GO:0003335);DNA metabolic process(GO:0006259);DNA catabolic process(GO:0006308) |
| **DNM2** | Dynamin-2 | G2/M transition of mitotic cell cycle(GO:0000086);mitochondrial fission(GO:0000266);G protein-coupled receptor internalization(GO:0002031);ventricular septum development(GO:0003281);regulation of transcription, DNA-templated(GO:0006355);post-Golgi vesicle-mediated transport(GO:0006892);Golgi to plasma membrane transport(GO:0006893);endocytosis(GO:0006897);receptor-mediated endocytosis(GO:0006898);phagocytosis(GO:0006909);signal transduction(GO:0007165);spermatogenesis(GO:0007283);response to light stimulus(GO:0009416);positive regulation of lamellipodium assembly(GO:0010592);synaptic vesicle budding from presynaptic endocytic zone membrane(GO:0016185);antigen processing and presentation of exogenous peptide antigen via MHC class II(GO:0019886);negative regulation of transforming growth factor beta receptor signaling pathway(GO:0030512);regulation of axon extension(GO:0030516);receptor internalization(GO:0031623);transferrin transport(GO:0033572);regulation of Rac protein signal transduction(GO:0035020);aorta development(GO:0035904);response to cocaine(GO:0042220);positive regulation of apoptotic process(GO:0043065);macropinocytosis(GO:0044351);positive regulation of nitric oxide biosynthetic process(GO:0045429);positive regulation of transcription, DNA-templated(GO:0045893);organelle fission(GO:0048285);synaptic vesicle transport(GO:0048489);neuron projection morphogenesis(GO:0048812);positive regulation of phagocytosis(GO:0050766);regulation of synapse structure or activity(GO:0050803);regulation of nitric-oxide synthase activity(GO:0050999);coronary vasculature development(GO:0060976);membrane organization(GO:0061024);membrane fusion(GO:0061025);cellular response to carbon monoxide(GO:0071245);cellular response to X-ray(GO:0071481);cellular response to nitric oxide(GO:0071732);postsynaptic neurotransmitter receptor internalization(GO:0098884);positive regulation of substrate adhesion-dependent cell spreading(GO:1900026);negative regulation of non-motile cilium assembly(GO:1902856);cellular response to dopamine(GO:1903351);regulation of Golgi organization(GO:1903358);positive regulation of sodium:potassium-exchanging ATPase activity(GO:1903408);negative regulation of membrane tubulation(GO:1903526);positive regulation of clathrin-dependent endocytosis(GO:2000370) |
| **DQX1** | ATP-dependent RNA helicase DQX1 | DNA duplex unwinding(GO:0032508) |
| **DSCAML1** | Down syndrome cell adhesion molecule-like protein 1 | cell fate determination(GO:0001709);cell adhesion(GO:0007155);homophilic cell adhesion via plasma membrane adhesion molecules(GO:0007156);axonogenesis(GO:0007409);axon guidance(GO:0007411);central nervous system development(GO:0007417);brain development(GO:0007420);dorsal/ventral pattern formation(GO:0009953);embryonic skeletal system morphogenesis(GO:0048704);dendrite self-avoidance(GO:0070593) |
| **DSCR4** | Down syndrome critical region protein 4 | biological_process(GO:0008150) |
| **DSCR8** | Down syndrome critical region protein 8 | biological_process(GO:0008150) |
| **DXO** | Decapping and exoribonuclease protein | mRNA catabolic process(GO:0006402);RNA destabilization(GO:0050779);nuclear mRNA surveillance(GO:0071028);nucleic acid phosphodiester bond hydrolysis(GO:0090305);NAD-cap decapping(GO:0110155) |
| **EEF1AKMT1** | EEF1A lysine methyltransferase 1 | protein methylation(GO:0006479);peptidyl-lysine methylation(GO:0018022) |
| **EFNA3** | Ephrin-A3;EFNA3 | cell-cell signaling(GO:0007267);axon guidance(GO:0007411);negative regulation of angiogenesis(GO:0016525);regulation of neuron differentiation(GO:0045664);ephrin receptor signaling pathway(GO:0048013);positive regulation of aspartic-type endopeptidase activity involved in amyloid precursor protein catabolic process(GO:1902961) |
| **F12** | Coagulation factor XII | plasma kallikrein-kinin cascade(GO:0002353);Factor XII activation(GO:0002542);proteolysis(GO:0006508);blood coagulation, intrinsic pathway(GO:0007597);positive regulation of plasminogen activation(GO:0010756);protein processing(GO:0016485);protein autoprocessing(GO:0016540);positive regulation of blood coagulation(GO:0030194);zymogen activation(GO:0031638);fibrinolysis(GO:0042730);innate immune response(GO:0045087);response to misfolded protein(GO:0051788);positive regulation of fibrinolysis(GO:0051919) |
| **FAM83H** | Protein FAM83H | positive regulation of cell migration(GO:0030335);biomineral tissue development(GO:0031214);protein localization to cytoskeleton(GO:0044380);intermediate filament cytoskeleton organization(GO:0045104) |
| **G6PC** | Glucose-6-phosphatase | glycogen metabolic process(GO:0005977);glycogen catabolic process(GO:0005980);gluconeogenesis(GO:0006094);triglyceride metabolic process(GO:0006641);steroid metabolic process(GO:0008202);response to carbohydrate(GO:0009743);regulation of gene expression(GO:0010468);glucose-6-phosphate transport(GO:0015760);response to food(GO:0032094);cellular response to insulin stimulus(GO:0032869);multicellular organism growth(GO:0035264);glucose homeostasis(GO:0042593);cholesterol homeostasis(GO:0042632);urate metabolic process(GO:0046415);phosphorylated carbohydrate dephosphorylation(GO:0046838);glucose 6-phosphate metabolic process(GO:0051156);response to resveratrol(GO:1904638) |

| | | |
|---|---|---|
| **GART** | Trifunctional purine biosynthetic protein adenosine-3 | brainstem development(GO:0003360);purine nucleotide biosynthetic process(GO:0006164);'de novo' IMP biosynthetic process(GO:0006189);glycine metabolic process(GO:0006544);purine nucleobase biosynthetic process(GO:0009113);purine ribonucleoside monophosphate biosynthetic process(GO:0009168);response to organic substance(GO:0010033);response to inorganic substance(GO:0010035);cerebellum development(GO:0021549);cerebral cortex development(GO:0021987);adenine biosynthetic process(GO:0046084);tetrahydrofolate biosynthetic process(GO:0046654) |
| **GIGYF1** | GRB10-interacting GYF protein 1 | biological_process(GO:0008150);insulin-like growth factor receptor signaling pathway(GO:0048009) |
| **GP6** | Platelet glycoprotein VI | enzyme linked receptor protein signaling pathway(GO:0007167);blood coagulation(GO:0007596);platelet activation(GO:0030168);leukocyte migration(GO:0050900) |
| **GPX5** | Epididymal secretory glutathione peroxidase | lipid metabolic process(GO:0006629);cellular response to oxidative stress(GO:0034599);hydrogen peroxide catabolic process(GO:0042744);oxidation-reduction process(GO:0055114);cellular oxidant detoxification(GO:0098869) |
| **GSTO2** | Glutathione S-transferase omega-2 | xenobiotic metabolic process(GO:0006805);L-ascorbic acid metabolic process(GO:0019852);oxidation-reduction process(GO:0055114);cellular response to arsenic-containing substance(GO:0071243);cellular oxidant detoxification(GO:0098869);glutathione derivative biosynthetic process(GO:1901687) |
| **HAGHL** | Hydroxyacylglutathione hydrolase-like protein | methylglyoxal catabolic process to D-lactate via S-lactoyl-glutathione(GO:0019243) |
| **HLCS** | Biotin protein ligase | biotin metabolic process(GO:0006768);protein biotinylation(GO:0009305);histone modification(GO:0016570);response to biotin(GO:0070781);histone biotinylation(GO:0071110) |
| **HMGN2** | Non-histone chromosomal protein HMG-17 | killing of cells of other organism(GO:0031640);antimicrobial humoral immune response mediated by antimicrobial peptide(GO:0061844) |
| **HSPA7** | Putative heat shock 70 kDa protein 7 | response to unfolded protein(GO:0006986);biological_process(GO:0008150);vesicle-mediated transport(GO:0016192);cellular response to unfolded protein(GO:0034620);protein refolding(GO:0042026);chaperone cofactor-dependent protein refolding(GO:0051085) |
| **HUNK** | Hormonally up-regulated neu tumor-associated kinase | protein phosphorylation(GO:0006468);signal transduction(GO:0007165);multicellular organism development(GO:0007275);intracellular signal transduction(GO:0035556) |
| **IFNAR2** | Interferon alpha/beta receptor 2 | cell surface receptor signaling pathway(GO:0007166);receptor signaling pathway via JAK-STAT(GO:0007259);response to virus(GO:0009615);cytokine-mediated signaling pathway(GO:0019221);response to interferon-alpha(GO:0035455);response to interferon-beta(GO:0035456);defense response to virus(GO:0051607);type I interferon signaling pathway(GO:0060337);regulation of type I interferon-mediated signaling pathway(GO:0060338) |
| **ITSN1** | Intersectin-1 | exocytosis(GO:0006887);endocytosis(GO:0006897);G protein-coupled receptor signaling pathway(GO:0007186);small GTPase mediated signal transduction(GO:0007264);brain development(GO:0007420);protein transport(GO:0015031);viral process(GO:0016032);positive regulation of kinase activity(GO:0033674);cellular protein localization(GO:0034613);positive regulation of apoptotic process(GO:0043065);negative regulation of neuron apoptotic process(GO:0043524);ephrin receptor signaling pathway(GO:0048013);synaptic vesicle endocytosis(GO:0048488);regulation of small GTPase mediated signal transduction(GO:0051056);positive regulation of protein kinase B signaling(GO:0051897);positive regulation of growth hormone secretion(GO:0060124);positive regulation of dendritic spine development(GO:0060999);membrane organization(GO:0061024);regulation of modification of postsynaptic actin cytoskeleton(GO:1905274);positive regulation of caveolin-mediated endocytosis(GO:2001288) |
| **JMJD8** | JmjC domain-containing protein 8 | regulation of glycolytic process(GO:0006110);positive regulation of I-kappaB kinase/NF-kappaB signaling(GO:0043123);regulation of pyruvate kinase activity(GO:1903302);positive regulation of sprouting angiogenesis(GO:1903672) |
| **KCNJ6** | G protein-activated inward rectifier potassium channel 2 | potassium ion transport(GO:0006813);regulation of ion transmembrane transport(GO:0034765);potassium ion import across plasma membrane(GO:1990573) |
| **KIAA1875** | WD repeat-containing protein 97 | |
| **KIF25** | Kinesin-like protein KIF25 | mitotic sister chromatid segregation(GO:0000070);organelle organization(GO:0006996);microtubule-based movement(GO:0007018);negative regulation of autophagy(GO:0010507);negative regulation of mitotic centrosome separation(GO:0046603);protein homotetramerization(GO:0051289);establishment of spindle orientation(GO:0051294);nucleus localization(GO:0051647) |
| **LMF1** | Lipase maturation factor 1 | triglyceride metabolic process(GO:0006641);endoplasmic reticulum to Golgi vesicle-mediated transport(GO:0006888);protein secretion(GO:0009306);protein glycosylation in Golgi(GO:0033578);chylomicron remnant clearance(GO:0034382);regulation of lipoprotein lipase activity(GO:0051004);positive regulation of lipoprotein lipase activity(GO:0051006);protein maturation(GO:0051604);regulation of cholesterol metabolic process(GO:0090181);regulation of triglyceride metabolic process(GO:0090207) |
| **LRPAP1** | Alpha-2-macroglobulin receptor-associated protein | negative regulation of receptor internalization(GO:0002091);negative regulation of very-low-density lipoprotein particle clearance(GO:0010916);negative regulation of protein binding(GO:0032091);regulation of receptor-mediated endocytosis(GO:0048259);negative regulation of cell death(GO:0060548);amyloid-beta clearance by transcytosis(GO:0150093);extracellular negative regulation of signal transduction(GO:1900116);negative regulation of amyloid-beta clearance(GO:1900222);positive regulation of amyloid-beta clearance(GO:1900223);negative regulation of signaling receptor activity(GO:2000272) |
| **LRRC14** | Leucine-rich repeat-containing protein 14 | negative regulation of NF-kappaB transcription factor activity(GO:0032088);negative regulation of toll-like receptor signaling pathway(GO:0034122) |
| **LRRC24** | Leucine-rich repeat-containing protein 24 | positive regulation of synapse assembly(GO:0051965) |
| **MEI1** | Meiosis inhibitor protein 1 | meiosis I(GO:0007127);male meiosis I(GO:0007141);spermatid development(GO:0007286);meiotic telomere clustering(GO:0045141) |
| **MGMT** | Methylated-DNA--protein-cysteine methyltransferase | DNA ligation(GO:0006266);DNA repair(GO:0006281);DNA methylation(GO:0006306);DNA dealkylation involved in DNA repair(GO:0006307);methylation(GO:0032259);cellular response to oxidative stress(GO:0034599);negative regulation of apoptotic process(GO:0043066);regulation of cysteine-type endopeptidase activity involved in apoptotic process(GO:0043281);response to ethanol(GO:0045471);positive regulation of DNA repair(GO:0045739);response to folic acid(GO:0051593);negative regulation of cell death(GO:0060548);mammary gland epithelial cell differentiation(GO:0060644);cellular response to organic cyclic compound(GO:0071407);cellular response to ionizing radiation(GO:0071479);positive regulation of double-strand break repair(GO:2000781) |
| **MZF1** | Myeloid zinc finger 1 | negative regulation of transcription by RNA polymerase II(GO:0000122);regulation of transcription, DNA-templated(GO:0006355);positive regulation of transcription by RNA polymerase II(GO:0045944) |
| **NME3** | Nucleoside diphosphate kinase 3 | purine nucleotide metabolic process(GO:0006163);nucleoside diphosphate phosphorylation(GO:0006165);GTP biosynthetic process(GO:0006183);pyrimidine nucleotide metabolic process(GO:0006220);UTP biosynthetic process(GO:0006228);CTP biosynthetic process(GO:0006241);apoptotic process(GO:0006915);nucleobase-containing small molecule interconversion(GO:0015949);regulation of apoptotic process(GO:0042981) |
| **NOXO1** | NADPH oxidase organizer 1 | superoxide metabolic process(GO:0006801);regulation of hydrogen peroxide metabolic process(GO:0010310);extracellular matrix disassembly(GO:0022617);positive regulation of catalytic activity(GO:0043085);regulation of respiratory burst(GO:0060263) |
| **NPAS4** | Neuronal PAS domain-containing protein 4 | regulation of transcription by RNA polymerase II(GO:0006357);learning(GO:0007612);short-term memory(GO:0007614);long-term memory(GO:0007616);cell differentiation(GO:0030154);regulation of synaptic transmission, GABAergic(GO:0032228);social behavior(GO:0035176);positive regulation of transcription by RNA polymerase II(GO:0045944);regulation of synaptic |

| | | |
|---|---|---|
| | | plasticity(GO:0048167);excitatory postsynaptic potential(GO:0060079);inhibitory postsynaptic potential(GO:0060080);cellular response to corticosterone stimulus(GO:0071386);inhibitory synapse assembly(GO:1904862) |
| **OLFM1** | Noelin | atrioventricular valve formation(GO:0003190);nervous system development(GO:0007399);positive regulation of gene expression(GO:0010628);negative regulation of gene expression(GO:0010629);positive regulation of epithelial to mesenchymal transition(GO:0010718);neuronal signal transduction(GO:0023041);regulation of axon extension(GO:0030516);positive regulation of apoptotic process(GO:0043065);cardiac epithelial to mesenchymal transition(GO:0060317) |
| **PFKL** | ATP-dependent 6-phosphofructokinase, liver type | fructose 6-phosphate metabolic process(GO:0006002);glucose catabolic process(GO:0006007);glycolytic process(GO:0006096);response to glucose(GO:0009749);fructose 1,6-bisphosphate metabolic process(GO:0030388);neutrophil degranulation(GO:0043312);negative regulation of insulin secretion(GO:0046676);protein homotetramerization(GO:0051289);canonical glycolysis(GO:0061621) |
| **PLEKHG5** | Pleckstrin homology domain-containing family G member 5 | G protein-coupled receptor signaling pathway(GO:0007186);endothelial cell chemotaxis(GO:0035767);positive regulation of apoptotic process(GO:0043065);positive regulation of I-kappaB kinase/NF-kappaB signaling(GO:0043123);regulation of small GTPase mediated signal transduction(GO:0051056);regulation of protein catabolic process at presynapse, modulating synaptic transmission(GO:0099575) |
| **PTPRS** | Receptor-type tyrosine-protein phosphatase S | protein dephosphorylation(GO:0006470);negative regulation of neuron projection development(GO:0010977);spinal cord development(GO:0021510);cerebellum development(GO:0021549);hippocampus development(GO:0021766);cerebral cortex development(GO:0021987);corpus callosum development(GO:0022038);negative regulation of axon extension(GO:0030517);negative regulation of interferon-alpha production(GO:0032687);negative regulation of interferon-beta production(GO:0032688);negative regulation of toll-like receptor 9 signaling pathway(GO:0034164);peptidyl-tyrosine dephosphorylation(GO:0035335);negative regulation of collateral sprouting(GO:0048671);negative regulation of axon regeneration(GO:0048681);modulation of chemical synaptic transmission(GO:0050804);negative regulation of dendritic spine development(GO:0061000);establishment of endothelial intestinal barrier(GO:0090557);regulation of postsynaptic density assembly(GO:0099151);synaptic membrane adhesion(GO:0099560) |
| **RASL10A** | Ras-like protein family member 10A | small GTPase mediated signal transduction(GO:0007264) |
| **RBM33** | RNA-binding protein 33 | |
| **RFX4** | Transcription factor RFX4 | regulation of transcription, DNA-templated(GO:0006355);regulation of transcription by RNA polymerase II(GO:0006357);telencephalon development(GO:0021537);negative regulation of smoothened signaling pathway involved in ventral spinal cord patterning(GO:0021914);positive regulation of transcription by RNA polymerase II(GO:0045944);cilium assembly(GO:0060271);regulation of protein processing(GO:0070613) |
| **RHBDL1** | Rhomboid-related protein 1 | proteolysis(GO:0006508);signal transduction(GO:0007165) |
| **RIBC2** | RIB43A-like with coiled-coils protein 2 | |
| **RRP1** | 60S acidic ribosomal protein P1 | nuclear-transcribed mRNA catabolic process, nonsense-mediated decay(GO:0000184);cytoplasmic translation(GO:0002181);rRNA processing(GO:0006364);translation(GO:0006412);translational initiation(GO:0006413);translational elongation(GO:0006414);regulation of translation(GO:0006417);SRP-dependent cotranslational protein targeting to membrane(GO:0006614);viral transcription(GO:0019083);activation of protein kinase activity(GO:0032147) |
| **RUNX1** | Runt-related transcription factor 1 | negative regulation of transcription by RNA polymerase II(GO:0000122);ossification(GO:0001503);regulation of cytokine-mediated signaling pathway(GO:0001959);chondrocyte differentiation(GO:0002062);regulation of transcription, DNA-templated(GO:0006355);regulation of transcription by RNA polymerase II(GO:0006357);negative regulation of gene expression(GO:0010629);hemopoiesis(GO:0030097);myeloid cell differentiation(GO:0030099);regulation of Wnt signaling pathway(GO:0030111);neuron differentiation(GO:0030182);negative regulation of granulocyte differentiation(GO:0030853);positive regulation of granulocyte differentiation(GO:0030854);positive regulation of interleukin-2 production(GO:0032743);regulation of intracellular estrogen receptor signaling pathway(GO:0033146);negative regulation of CD4-positive, alpha-beta T cell differentiation(GO:0043371);positive regulation of CD8-positive, alpha-beta T cell differentiation(GO:0043378);regulation of regulatory T cell differentiation(GO:0045589);regulation of cell differentiation(GO:0045595);regulation of keratinocyte differentiation(GO:0045616);regulation of myeloid cell differentiation(GO:0045637);regulation of megakaryocyte differentiation(GO:0045652);positive regulation of angiogenesis(GO:0045766);positive regulation of transcription, DNA-templated(GO:0045893);positive regulation of transcription by RNA polymerase II(GO:0045944);peripheral nervous system neuron development(GO:0048935);regulation of B cell receptor signaling pathway(GO:0050855);hematopoietic stem cell proliferation(GO:0071425);regulation of hematopoietic stem cell differentiation(GO:1902036);regulation of bicellular tight junction assembly(GO:2000810) |
| **RYR1** | Ryanodine receptor 1 | response to hypoxia(GO:0001666);outflow tract morphogenesis(GO:0003151);calcium ion transport(GO:0006816);muscle contraction(GO:0006936);release of sequestered calcium ion into cytosol by sarcoplasmic reticulum(GO:0014808);response to caffeine(GO:0031000);ion transmembrane transport(GO:0034220);skin development(GO:0043588);ossification involved in bone maturation(GO:0043931);skeletal muscle fiber development(GO:0048741);release of sequestered calcium ion into cytosol(GO:0051209);protein homotetramerization(GO:0051289);regulation of cytosolic calcium ion concentration(GO:0051480);cellular response to calcium ion(GO:0071277);cellular response to caffeine(GO:0071313);regulation of cardiac conduction(GO:1903779) |
| **SERPINH1** | Serpin H1 | chondrocyte development involved in endochondral bone morphogenesis(GO:0003433);response to unfolded protein(GO:0006986);negative regulation of endopeptidase activity(GO:0010951);collagen fibril organization(GO:0030199);collagen biosynthetic process(GO:0032964);protein maturation(GO:0051604) |
| **SIRPD** | Signal-regulatory protein delta | |
| **SLC16A1** | Monocarboxylate transporter 1 | pyruvate metabolic process(GO:0006090);lipid metabolic process(GO:0006629);centrosome cycle(GO:0007098);monocarboxylic acid transport(GO:0015718);mevalonate transport(GO:0015728);response to food(GO:0032094);lactate transmembrane transport(GO:0035873);plasma membrane lactate transport(GO:0035879);glucose homeostasis(GO:0042593);regulation of insulin secretion(GO:0050796);leukocyte migration(GO:0050900);behavioral response to nutrient(GO:0051780);cellular response to organic cyclic compound(GO:0071407) |
| **SLC35C1** | GDP-fucose transporter 1 | carbohydrate transport(GO:0008643);lipid glycosylation(GO:0030259);protein O-linked fucosylation(GO:0036066);GDP-fucose import into Golgi lumen(GO:0036085);negative regulation of Notch signaling pathway(GO:0045746) |
| **SLC5A2** | Sodium/glucose cotransporter 2 | carbohydrate metabolic process(GO:0005975);sodium ion transport(GO:0006814);hexose transmembrane transport(GO:0008645);transmembrane transport(GO:0055085);glucose transmembrane transport(GO:1904659) |
| **SMC1B** | Structural maintenance of chromosomes protein 1B | sister chromatid cohesion(GO:0007062);meiotic cell cycle(GO:0051321) |
| **SON** | Protein SON | microtubule cytoskeleton organization(GO:0000226);mitotic cytokinesis(GO:0000281);mRNA processing(GO:0006397);RNA splicing(GO:0008380);negative regulation of apoptotic process(GO:0043066);regulation of RNA splicing(GO:0043484);regulation of mRNA splicing, via spliceosome(GO:0048024);regulation of cell cycle(GO:0051726) |
| **STK18** | Serine/threonine-protein kinase PLK4 | G2/M transition of mitotic cell cycle(GO:0000086);mitotic prometaphase(GO:0000236);mitotic cell cycle(GO:0000278);protein phosphorylation(GO:0006468);centriole replication(GO:0007099);regulation of G2/M transition of mitotic cell cycle(GO:0010389);regulation of cytokinesis(GO:0032465);positive regulation of centriole replication(GO:0046601);trophoblast giant cell differentiation(GO:0060707);ciliary basal body-plasma membrane docking(GO:0097711);de novo centriole assembly involved in multi-ciliated epithelial cell differentiation(GO:0098535) |
| **STK19** | Serine/threonine-protein kinase 19 | protein phosphorylation(GO:0006468);phosphorylation(GO:0016310);positive regulation of Ras protein signal transduction(GO:0046579) |

| | | |
|---|---|---|
| **SUMO3** | Small ubiquitin-related modifier 3 | protein sumoylation(GO:0016925);negative regulation of DNA binding(GO:0043392) |
| **TAPBP** | Tapasin | MHC class I protein complex assembly(GO:0002397);MHC class Ib protein complex assembly(GO:0002398);antigen processing and presentation of peptide antigen via MHC class I(GO:0002474);antigen processing and presentation of exogenous peptide antigen via MHC class I, TAP-dependent(GO:0002479);retrograde vesicle-mediated transport, Golgi to endoplasmic reticulum(GO:0006890);immune response(GO:0006955);regulation of gene expression(GO:0010468);peptide transport(GO:0015833);antigen processing and presentation of endogenous peptide antigen via MHC class I(GO:0019885);peptide antigen stabilization(GO:0050823);regulation of protein complex stability(GO:0061635);protein-containing complex assembly(GO:0065003);vesicle fusion with endoplasmic reticulum-Golgi intermediate compartment (ERGIC) membrane(GO:1990668) |
| **TDH** | Inactive L-threonine 3-dehydrogenase, mitochondrial | threonine catabolic process(GO:0006567);L-threonine catabolic process to glycine(GO:0019518);oxidation-reduction process(GO:0055114) |
| **TDH** | Serine dehydratase-like | L-serine catabolic process(GO:0006565);threonine catabolic process(GO:0006567);biological_process(GO:0008150);L-threonine catabolic process to glycine(GO:0019518) |
| **TDH** | L-serine dehydratase/L-threonine deaminase | gluconeogenesis(GO:0006094);cellular amino acid metabolic process(GO:0006520);L-serine catabolic process(GO:0006565);threonine catabolic process(GO:0006567);L-threonine catabolic process to glycine(GO:0019518);pyruvate biosynthetic process(GO:0042866) |
| **TYMP** | Thymidine phosphorylase | mitochondrial genome maintenance(GO:0000002);angiogenesis(GO:0001525);pyrimidine nucleobase metabolic process(GO:0006206);pyrimidine nucleoside metabolic process(GO:0006213);chemotaxis(GO:0006935);signal transduction(GO:0007165);cell differentiation(GO:0030154);regulation of myelination(GO:0031641);pyrimidine nucleoside salvage(GO:0043097);pyrimidine nucleoside catabolic process(GO:0046135);regulation of transmission of nerve impulse(GO:0051969);regulation of gastric motility(GO:1905333) |
| **UNC45A** | Protein unc-45 homolog A | muscle organ development(GO:0007517);cell differentiation(GO:0030154);chaperone-mediated protein folding(GO:0061077) |
| **USP50** | Inactive ubiquitin carboxyl-terminal hydrolase 50 | ubiquitin-dependent protein catabolic process(GO:0006511);endosome organization(GO:0007032);Ras protein signal transduction(GO:0007265);protein deubiquitination(GO:0016579);positive regulation of interleukin-18 production(GO:0032741);nuclear speck organization(GO:0035063);positive regulation of interleukin-1 beta secretion(GO:0050718);protein K63-linked deubiquitination(GO:0070536);protein K48-linked deubiquitination(GO:0071108);positive regulation of NLRP3 inflammasome complex assembly(GO:1900227);positive regulation of cysteine-type endopeptidase activity(GO:2001056) |
| **USP8** | Ubiquitin carboxyl-terminal hydrolase 8 | mitotic cytokinesis(GO:0000281);ubiquitin-dependent protein catabolic process(GO:0006511);endosome organization(GO:0007032);Ras protein signal transduction(GO:0007265);protein deubiquitination(GO:0016579);regulation of protein stability(GO:0031647);regulation of protein localization(GO:0032880);protein K63-linked deubiquitination(GO:0070536);protein K48-linked deubiquitination(GO:0071108);cellular response to dexamethasone stimulus(GO:0071549);positive regulation of canonical Wnt signaling pathway(GO:0090263);regulation of protein catabolic process at postsynapse, modulating synaptic transmission(GO:0099576);cellular response to nerve growth factor stimulus(GO:1990090) |
| **VPS37B** | Vacuolar protein sorting-associated protein 37B | protein targeting to membrane(GO:0006612);protein targeting to vacuole(GO:0006623);protein transport(GO:0015031);endosomal transport(GO:0016197);macroautophagy(GO:0016236);viral life cycle(GO:0019058);endosome transport via multivesicular body sorting pathway(GO:0032509);multivesicular body assembly(GO:0036258);viral budding via host ESCRT complex(GO:0039702);ubiquitin-dependent protein catabolic process via the multivesicular body sorting pathway(GO:0043162);intracellular transport of virus(GO:0075733);positive regulation of viral release from host cell(GO:1902188);positive regulation of viral budding via host ESCRT complex(GO:1903774) |
| **WRB** | Tail-anchored protein insertion receptor WRB | posttranslational protein targeting to endoplasmic reticulum membrane(GO:0006620);sensory perception of sound(GO:0007605);synapse organization(GO:0050808);otic vesicle development(GO:0071599);tail-anchored membrane protein insertion into ER membrane(GO:0071816) |
| **YBX2** | Y-box-binding protein 2 | transcription by RNA polymerase II(GO:0006366);spermatogenesis(GO:0007283);translational attenuation(GO:0009386);oocyte development(GO:0048599);positive regulation of cold-induced thermogenesis(GO:0120162) |
| **ZBTB22** | Zinc finger and BTB domain-containing protein 22 | regulation of transcription by RNA polymerase II(GO:0006357) |
| **ZNF497** | Zinc finger protein 497 | negative regulation of transcription by RNA polymerase II(GO:0000122);positive regulation of transcription by RNA polymerase II(GO:0045944) |
| **ZNF837** | Zinc finger protein 837 | |

**Supplementary Table 3.** *RUNX1* mRNA expression and DNA methylation levels in myocardial samples from donors with and without DS

| Sample_ID | Sample_Group | Gender | Age (years) | β-value (*RUNX1*) | *RUNX1* relative fold expression |
|---|---|---|---|---|---|
| **Heart_1** | NDS | M | 57 | 0.079 | 2.71 |
| **Heart_5** | NDS | M | 61 | 0.119 | 3.17 |
| **Heart_9** | NDS | M | 70 | 0.043 | 1.30 |
| **Heart_14** | NDS | M | 80 | 0.321 | 1.12 |
| **Heart_21** | NDS | F | 47 | 0.041 | 1.49 |
| **Heart_24** | NDS | M | 69 | 0.186 | 1.59 |
| **Heart_53** | NDS | F | 38 | 0.1 | 0.32 |
| **Heart_54** | NDS | M | 47 | 0.073 | 0.38 |
| **Heart_3** | NDS | F | 55 | ND | 0.80 |
| **Heart_4** | NDS | F | 63 | ND | 0.33 |
| **Heart_7** | NDS | F | 74 | ND | 0.19 |
| **Heart_8** | NDS | F | 69 | ND | 0.60 |
| **Heart_10** | NDS | F | 67 | ND | 0.17 |
| **Heart_25** | NDS | F | 66 | ND | 0.56 |
| **Heart_5824** | NDS | M | 91 | ND | 1.02 |
| **Heart_5863** | NDS | F | 62 | ND | 0.24 |
| **Heart_18** | DS | M | 58 | 0.302 | 12.80 |
| **Heart_17** | DS | M | 63 | 0.294 | 6.59 |
| **Heart _15** | DS | M | 72 | 0.291 | 1.06 |
| **Heart_37** | DS | M | 22 | 0.314 | 5.82 |
| **Heart_48** | DS | F | 48 | 0.331 | 1.61 |
| **Heart_23** | DS | M | 59 | 0.333 | 4.57 |
| **Heart_39** | DS | F | 1 | 0.396 | 4.25 |
| **Heart_19** | DS | M | 39 | 0.295 | 22.71 |
| **Heart_41** | DS | M | 64 | 0.313 | 3.57 |
| **Heart_33** | DS | M | 40 | 0.315 | 19.50 |
| **Heart_38** | DS | M | 1 | 0.313 | 3.46 |
| **Heart_40** | DS | M | 10 | 0.321 | 3.85 |
| **Heart_49** | DS | F | 31 | ND | 3.25 |
| **Heart_55** | DS | F | 51 | ND | 0.68 |
| **Heart_8171** | DS | F | 56 | ND | 1.98 |
| **Heart_8925** | DS | F | 33 | ND | 11.91 |

ND: No β-value data available
